# Synchronization through uncorrelated noise in excitatory-inhibitory networks

**DOI:** 10.1101/2021.10.29.466430

**Authors:** Rebscher Lucas, Obermayer Klaus, Metzner Christoph

## Abstract

Gamma rhythms play a major role in many different processes in the brain, such as attention, working memory and sensory processing. While typically considered detrimental, counterintuitively noise can sometimes have beneficial effects on communication and information transfer. Recently, Meng and Riecke showed that synchronization of interacting networks of inhibitory neurons in the gamma band increases while synchronization within these networks decreases when neurons are subject to uncorrelated noise. However, experimental and modeling studies point towards an important role of the pyramidal-interneuronal network gamma (PING) mechanism in the cortex. Therefore, we investigated the effect of uncorrelated noise on the communication between excitatory-inhibitory networks producing gamma oscillations via a PING mechanism. Our results suggest that synaptic noise can have a supporting role in facilitating inter-regional communication and that noise-induced synchronization between networks is generated via a different mechanism than when synchronization is mediated by strong synaptic coupling. Noise-induced synchronization is achieved by lowering synchronization within networks which allows the respective other network to impose its own gamma rhythm resulting in synchronization between networks.

## Introduction

Synchronous oscillatory activity in high and low frequency ranges has been proposed to underlie coordinated communication between distributed neural systems [1–4]. Especially gamma rhythms (high-frequency oscillations in the 30-90Hz range) have been studied extensively and have been related to perception [5], attention [6], memory [7], consciousness [8] and synaptic plasticity [9]. Furthermore, pathological brain states in neurological and psychiatric disorders, such as Alzheimer’s, autism and schizophrenia, have been linked to dysfunctional neural oscillations in the gamma band [10, 11].

Mechanistically, *in vitro* and *in vivo* gamma rhythms are mainly produced by two mechanisms termed interneuron network gamma (ING) and pyramidal-interneuron network gamma (PING) [12–14]. The ING mechanism is based on the mutual inhibition of inhibitory neurons, which act as a gate that temporarily suppresses firing until inhibition wears off and the neurons fire in increased synchrony [15]. On the other hand, the PING mechanism is based on the interplay between excitation and inhibition [15]. Firing of excitatory neurons prompts firing of inhibitory neurons which in turn temporarily suppress further firing, ultimately leading to coherent activity in both groups.

Gamma rhythms have been proposed to underlie efficient communication between different brain regions [16–18]. For example, the communication-through-coherence (CTC) hypothesis [17, 19] posits that synchronization of two brain regions or circuits in the gamma band allows for a more efficient transfer of information between them. Over the last years, this proposal has been supported by considerable experimental evidence, such as, gain modulation of both neural and behavioural responses in the gamma band [20], attentional enhancement of gamma-band synchrony between neural populations [21, 22], and covariations in transfer entropy and gamma-band synchrony [23, 24]. Naturally, neural regions and their communication are subject to various sources of noise and typically noise is assumed to be detrimental to the quality of the signal transfer between the regions and their synchronization. Computational models have for example confirmed that noise can reduce the synchronization of excitatory-inhibitory (EI) networks [25, 26]. However, in non-linear biological systems one can also observe a helpful role of noise under certain conditions [27, 28], such as for example stochastic resonance, the noise-induced improved response to a weak signal, has been well documented experimentally and theoretically in several neural systems [29–31].

In a recent computational study, Meng and Riecke [32] demonstrated that noise can synchronize poulation rhythms generated by individual oscillator networks. They showed that noise induced synchronization despite the noise input between different oscillator networks being uncorrelated. This between-network synchronization emerges as the uncorrelated noise introduces heterogeneity within the networks, thereby weakening intra-network synchronization, and thus allowing for the second network to control a substantial fraction of the network activity. While they demonstrate that this type of synchronization emerges in different settings, their findings remain restricted to networks coupled by inhbition, i.e. they do not investigate the synchronization of PING networks.

Therefore, in this study, we model two interconnected excitatory-inhibitory networks, producing gamma oscillations through a PING mechanism, in different network settings and analyze how synchronization within and across the networks changes depending on the strength of uncorrelated noise input to the networks. Our results extend the findings from Meng and Riecke [32] and suggest that uncorrelated noise can also have a supporting role in facilitating inter-regional communication in PING networks.

Importantly, our models can be used as a basis to investigate mechanistic explanations for altered neuronal dynamics in psychiatric disorders, since for example, disturbances in neuronal oscillations in the gamma band, especially reduced synchronization, are a key finding in schizophrenia [11].

## Materials and methods

We first replicated the results of Meng and Riecke [32] using two interacting inhibitory networks producing gamma rhythms via an ING mechanism where each neuron was subject to independent noise. Once we were able to replicate their results in our model implementation, we proceeded to the next step and extended the model to two interacting excitatory-inhibitory networks showing a PING mechanism. In this case, we looked at two variants: the simpler case of all-to-all connectivity and the biologically more plausible case of sparse random connectivity. Conclusively, in the following we define the three scenarios that we simulated, evaluated and compared:

- **Scenario 1**: In this scenario we replicated the case of two interacting inhibitory networks with all-to-all connectivity investigated by Meng and Riecke [32]. Importantly, each neuron was subject to uncorrelated noise and both networks displayed rhythmic gamma band activity produced by the ING mechanism.
- **Scenario 2**: This scenario extended scenario 1 to two all-to-all coupled excitatory-inhibitory networks driven by the PING mechanism. It also built the foundation for scenario 3 and potentially more complex variants used in future work.
- **Scenario 3**: In the last scenario we moved from all-to-all connectivity to random sparse connectivity to assess to which degree independent noise would also be able to have a beneficial effect in networks with a biologically more plausible connectivity.

### Network Models

In scenario 1, we considered a model of two coupled inhibitory networks analoguous to Meng and Riecke [32]. Both populations consisted of 1000 inhibitory neurons and received input in form of random uncorrelated spike trains of 800 excitatory neurons according to a Poisson schedule. Each population had recurrent connections and both networks were bidirectionally coupled (see Fig 1A for a schematic of the ING network setup). As connectivity in scenario 1 is all-to-all, the connection probability *p* was 1.0 for all connection types.

**Fig 1.**
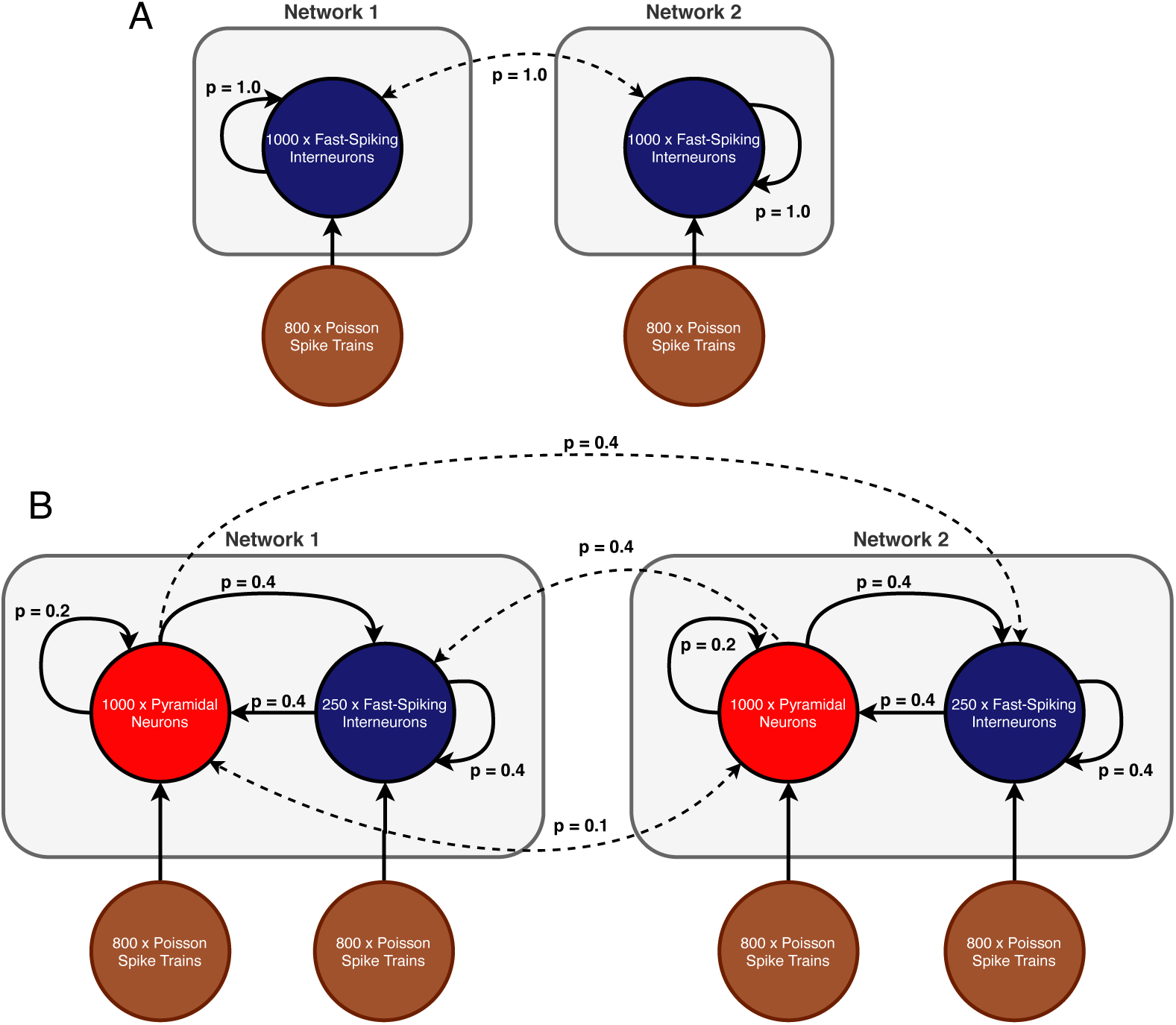
Network models. (A) **Scenario 1:** Two all-to-all coupled networks, each consisting of an inhibitory population. In both populations all neurons receive independent noise and synaptic coupling is stronger within networks than across networks. (B) **Scenario 2 and 3:** Two coupled excitatory-inhibitory networks in which each population is again subject to uncorrelated noise. In comparison to scenario 1, inter-network communication is mediated exclusively by excitatory connections. The displayed connection probabilities are used in the sparse random network in scenario 3. In case of scenario 2, probability *p* is instead set to 1.0 for all connection pairs.

In scenarios 2 and 3, each network consisted of a population of 250 inhibitory neurons as well as a population of 1000 excitatory neurons (See Fig 1B for a schematic of the setup). The difference between the two scenarios lies in the particular connectivity scheme (all-to-all vs sparse random) and the synaptic coupling strengths. Excitatory and inhibitory populations within each network were recurrently connected. However, inter-network communication was mediated solely by excitatory synapses that target both the inhibitory and excitatory populations in the respective other network. The connection probabilities were configured according to the specific scenario and are shown in Table 1. While all populations were again subject to uncorrelated noise, the inhibitory populations received a lower proportion of noise to avoid that the spiking of inhibitory neurons became dominated by the noise input, since this would have caused the PING rhythms to collapse [33].

**Table 1.**
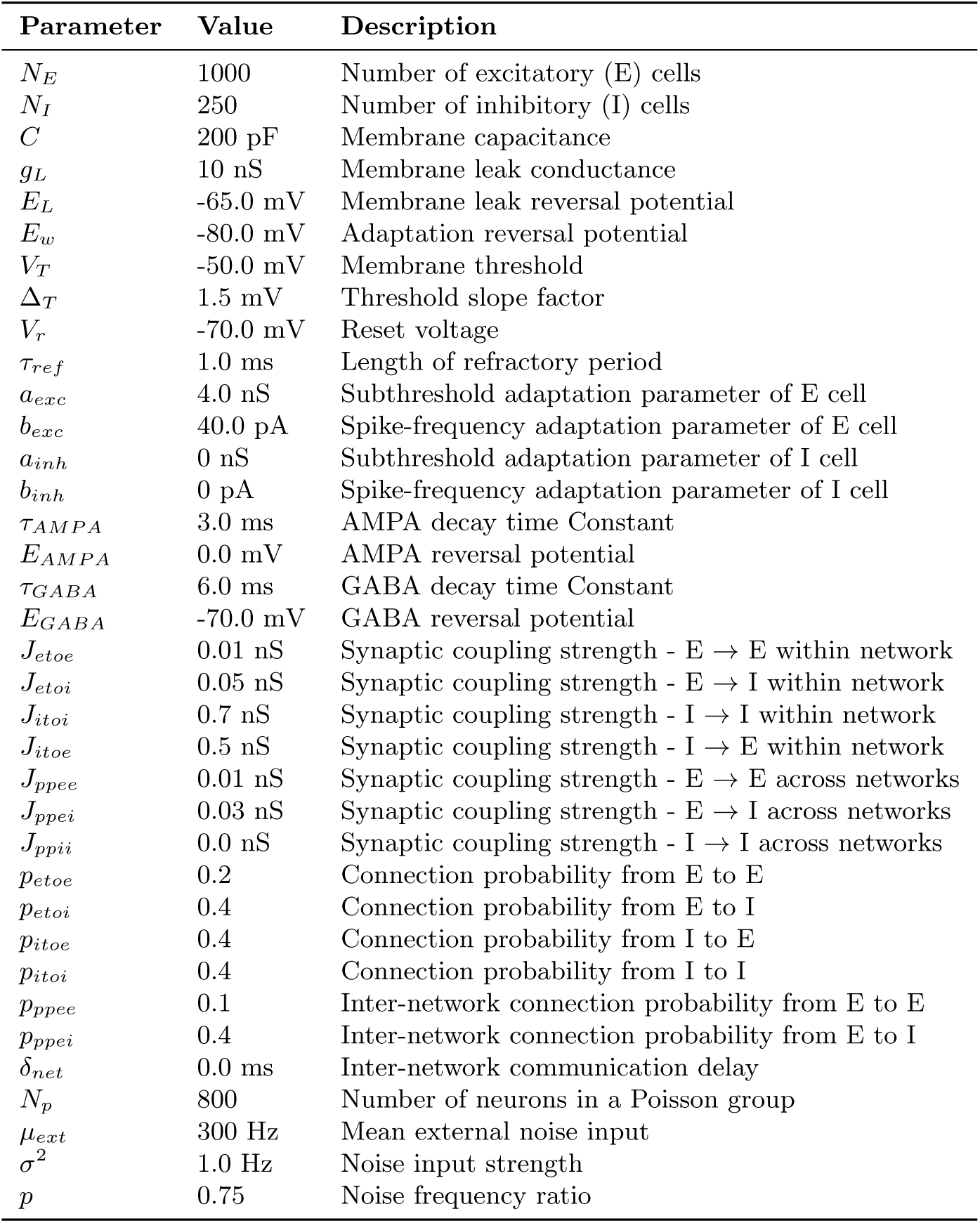
A list of all relevant model parameters, including parameters of AdEx model, GABA- and AMPA-mediated synapses, synaptic noise and connection probabilities. Parameters such as coupling strengths and connection probabilities vary across scenarios.

### Neuron Model

As a neuron model we used the adaptive exponential integrate-and-fire (AdEx) model proposed by Brette and Gerstner [34] and its membrane potential time course is described in Eq 1. To improve readability, we extracted the internal membrane dynamics to the current *I*_*ion*_ in Eq 2. Adaptation is modeled by the parameter *w* as defined in Eq 3.

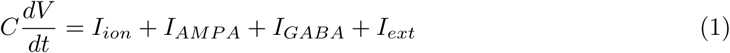

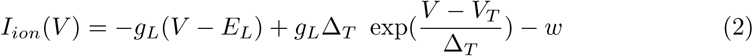

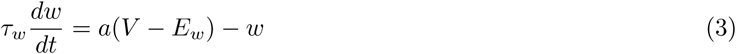

where *C* is the membrane capacitance, *E*_*L*_ the leak reversal potential, *E*_*w*_ the adaptation reversal potential, *V* the membrane voltage at time step *t, V*_*T*_ the membrane threshold, *V*_*reset*_ the reset potential, Δ*T* the slope factor, *a* the adaptation coupling parameter, *τ*_*w*_ the adaptation time constant and *g*_*L*_ the leak conductance. When the membrane potential exceeds the spiking threshold, the voltage is reset to *V*_*r*_ and clamped for a refractory time *T*_*ref*_. Furthermore, the spike-triggered adaptation increment *b* is added to the adaptation current. Each neuron received post-synaptic currents *I*_*GABA*_ and *I*_*AMP A*_ which we further specify below. Finally, the current *I*_*ext*_ represents the external noise input that we define below.

### Synapse Model

Synaptic currents for GABAergic as well as glutamatergic (AMPA) synapses are defined in Eq 4 and Eq 6, respectively. We used a conductance-based model and the respective synaptic conductance is modeled as a dynamic variable with an instantaneous rise on each pre-synaptic spike and an exponential decay over time. The exponential decay is defined in Eq 5 and Eq 7 for AMPA and GABA, respectively.

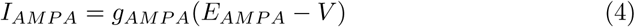

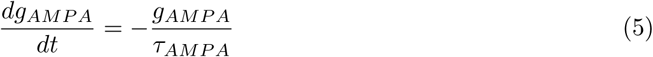

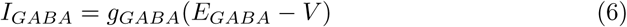

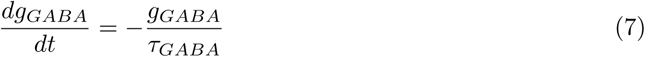

where *g*_*GABA*_, *g*_*AMP A*_ are the dynamic synaptic conductances, *E*_*AMP A*_ and *E*_*GABA*_ the characteristic reversal potentials, *V* the membrane potential of the post-synaptic neuron and lastly *τ*_*AMP A*_, *τ*_*GABA*_ represent the time constants that define the speed of the exponential decay.

The synaptic conductances *g*_*GABA*_ and *g*_*AMP A*_ is modeled for each neuron and their instantaneous rise is described by the following update rules executed on every pre-synaptic spike:

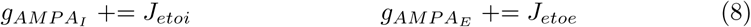

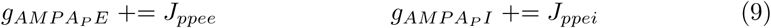

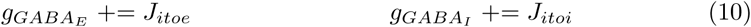

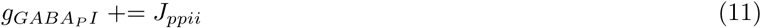

where *J*_*etoi*_, *J*_*etoe*_, *J*_*itoe*_, *J*_*itoi*_ are the synaptic coupling strengths between excitatory or inhibitory neurons *within* a network, whereas *J*_*ppee*_, *J*_*ppei*_, *J*_*ppii*_ are the coupling strengths *across* networks. We deliberately left out the coupling strength *J*_*ppie*_ of synapses originating from inhibitory neurons in one network and targeting excitatory neurons in another network, since this type of connection is not present in any of our scenarios. Further, note that *J*_*ppii*_ was only present in the inhibitory networks of scenario 1 whereas *J*_*ppee*_, *J*_*ppei*_ were only present in the excitatory-inhibitory networks of scenarios 2 and 3.

### Noise Model

We adopted the noise framework that Meng and Riecke [32] used to generate uncorrelated noise for each neuron. Each population in every scenario was subject to substantial synaptic noise in form of a group of *N*_*p*_ = 800 Poisson neurons. We used the mean input *μ*, the input strength *σ*^2^ and the frequency ratio *p* as free parameters in our explorations.

For each Poisson neuron we generated spike trains according to a Poisson process with a rate *λ* that was dependent on the free parameters *μ* and *σ*^2^, as shown in Eq 12, where *N*_*p*_ (e.g., *N*_*p*_ = 800) is the number of Poisson neurons.

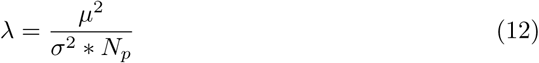

Importantly, the rate differed between the two networks and network 2 received a lower rate than network 1. This ultimately determined the natural frequency of the network activity and was performed to set the networks to a desynchronized state. Therefore, the difference in the noise input between the two networks lay only in the rate *λ*, the strength of a Poisson generated spike stayed the same within and across networks. The rate difference between network 1 and network 2 was controlled by the frequency ratio *p* with *p* ∈ [0, 1] as follows

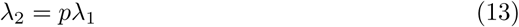

Besides generating spike trains, we also needed to model the impact of a generated pre-synaptic spike on a post-synaptic neuron. This was modeled in form of an external current *I*_*ext*_ with

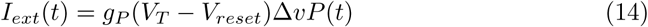

The conductance *g*_*P*_ was included to ensure correct physical units and we set it to a constant value of 1 *nS. P* (*t*) is the number of pre-synaptic Poisson spikes arriving at time step *t* at the respective post-synaptic neuron. The difference between the membrane threshold *V*_*T*_ and the reset potential *V*_*reset*_ of the respective post-synaptic neuron is precisely the amount of voltage needed to produce a spike independent of its current membrane potential. However, the extent of this driving force is controlled by a dimensionless input strength Δ*v*. Analog to the rate *λ*, Δ*v* is also dependent on the free parameters *σ*^2^ and *μ* according to

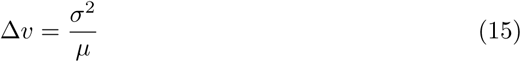

We can observe based on Eq 12 and Eq 15 that an increase in the noise strength *σ*^2^ leads on one hand to an increase in the spike strength Δ*v*_*i*_ and on the other hand to a decrease in the rate *λ*.

### Parameter Explorations

Two-dimensional explorations were visualized in form of heat maps where the colors were mapped to the value of a specific measure. We decided to use three different measures which are presented in the following.

#### Phase Synchronization between two Network

The synchronization between the networks is perfect if the phase difference over time is constant as we might have considerable delay regulated by parameter *δ*_*net*_. Thus, the synchronization between networks was quantified by the mean phase coherence of the networks’ activity. We used the average *n*_*i*_ of the neurons’ voltage traces inside a network as a surrogate for the network level activity

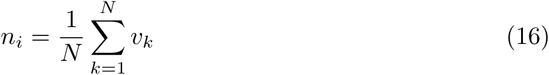

where the sum is over all *N* neurons inside network *i* and *v*_*k*_ is the recorded voltage trace of neuron *k*.

Before extracting the phase information, we bandpass filtered the signal *n*_*i*_ to ensure that the extracted phase information is meaningful. As a filter we used a second-order Butterworth filter with a lowcut frequency of 30 Hz and a highcut frequency of 120 Hz implemented in the *scipy*^1^ package `scipy.signal.filter_design.butter`. Next, we extracted the phases *φ*_*i*_ from the filtered signal by applying the Hilbert transform implemented in the *scipy* package under `scipy.signal.hilbert.` Finally, the mean phase coherence between the two networks was calculated as

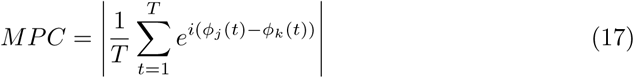

where *φ*_*j*_(*t*) is the phase of the signal of population *j* at time *t* and *T* the total number of time steps. If the phase difference between two signals is constant over time *MPC* = 1. In this case, the signals are said to be *phase locked*. On the other hand, *MPC* = 0 indicates no correlation between the phases.

#### Phase Synchronization within a Population

When determining the phase synchronization within a population, the population activity was again represented as the average of the voltage traces of the corresponding neurons. Further, the same preprocessing steps as above were applied. However, instead of using the mean phase coherence, we chose the Kuramoto order parameter because perfect synchronization inside a homogeneous population is expected to be equivalent to zero phase lag synchronization. The Kuramoto order parameter was then calculated according to

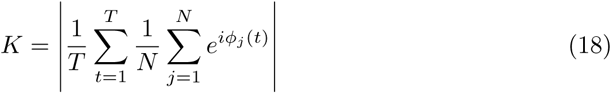

where *φ*_*j*_(*t*) is now the phase of signal of an individual neuron *j* at time *t, T* the total number of time steps and *N* the total number of neurons in the population.

In contrast to MPC which measures the *consistency* of phase *differences*, the Kuramoto order parameter is based on the divergence between the phases. Thus, perfect synchronization of *K* = 1 is reached if the signals are identical which implies a zero phase lag between the signals. In the worst case, the signals are anti-phase synchronized, resulting in *K* = 0. Importantly, while MPC and Kuramoto order parameter are both labeled as *phase synchronization* measures, they measure distinct forms of synchronization. If we assume two signals that display perfect anti-phase synchronization, the Kuramoto order parameter would be 0 as the phase lag is maximal over time, while the MPC would be 1 as the phase differences are constant over time.

#### Frequency Locking of two Networks

The difference in the frequencies of two oscillators is a measure of frequency locking (also sometimes referred to as *entrainment*) [35]. To quantify frequency locking between two networks, we used the same measure as Meng and Riecke [32], namely the ratio between the dominant frequencies of the networks. First, the network signals were transformed to the frequency-domain by applying Welch’s method [36] offered by the Python package *matplotlib*^2^ in the module `matplotlib.mlab.psd`. Next, the dominant frequency *f*_*i*_ of each of the two networks was determined by extracting the frequency with the highest power. Finally, the ratio between the two extracted frequencies was calculated as follows

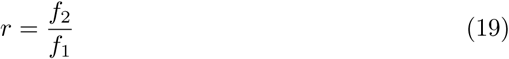

with *f*_2_ ≤ *f*_1_ and *f*_1_, *f*_2_ *>* 0. The larger dominant frequency corresponds to *f*_1_ and the lower one to *f*_2_ to ensure that *r* ∈ (0, 1]. A ratio of 1 indicates perfect frequency locking as the two frequencies equal whereas values close to 0 indicate maximal mismatch between the two frequencies. Noteworthy, in our simulations, network 1 always had a greater or equal dominant frequency than network 2 because it received stronger noise input.

### Code and Simulations

The source code is publicly accessible on GitHub^3^. All code was written in Python 3.7. We used the spiking neural simulator *Brian 2* [37] *(2*.*3*.*0*.*2)* for model simulations. To run large scale parameter explorations we used the exploration library *mopet* [38] *(0*.*1*.*3)*. All plots were created with the Python library *matplotlib*^4^ *(3*.*1*.*2)* and signal processing steps were performed with the *scipy* ^5^ *(1*.*4*.*1)* package.

To ensure that our simulation results were robust and could be reproduced in a different simulation setting, we evaluated our results with different step sizes (1.0 ms, 0.5 ms, 0.05 ms), simulation durations (0.5 s, 1 s, 3 s, 5 s) and integration methods. Regarding the integration methods, we used three different methods implemented in *Brian 2* ^6^: *heun* (stochastic Heun method), *euler* (forward Euler integration) and *milstein* (derivative-free Milstein method). The results did not vary noticeably in all cases. For our final simulations, we used a step size of 0.05, a duration of 5*s* and the *heun* integration method.

## Results

### Uncorrelated noise can synchronize networks of ING populations

In scenario 1, we first investigated the effect of uncorrelated noise input on the synchronization of two coupled inhibitory networks that produce a gamma oscillation via an ING mechanism (similar to Meng and Riecke [32]).

In this scenario, both inhibitory networks were driven by external, uncorrelated noise. Importantly, network 1 had a higher natural frequency than network 2 as network 1 was subject to stronger noise. This difference was controlled by the frequency ratio parameter *p*. We then explored whether, and if so to what extent, certain noise strengths *σ*^2^ might be able to enhance the synchronization between the networks in relation to the frequency ratio *p*.

#### Exploration

For a fixed noise ratio *p*, e.g., *p* = 0.85, we observed that the dominant frequency ratio and the phase synchronization between networks improved if the noise strength *σ*^2^ was sufficiently increased (Fig 2A). At the same time, the within phase synchronization of the respective inhibitory networks decreased. For sufficiently large frequency ratios *p* (typically *p >* 0.75), an increase above a certain threshold in the noise strength generally improved the synchronization between the networks initially (Fig 2A). Additionally, higher noise frequency ratios seemed to lower the minimal noise strength *σ*^2^ required to transition the two networks to a frequency and phase locked state.

**Fig 2.**
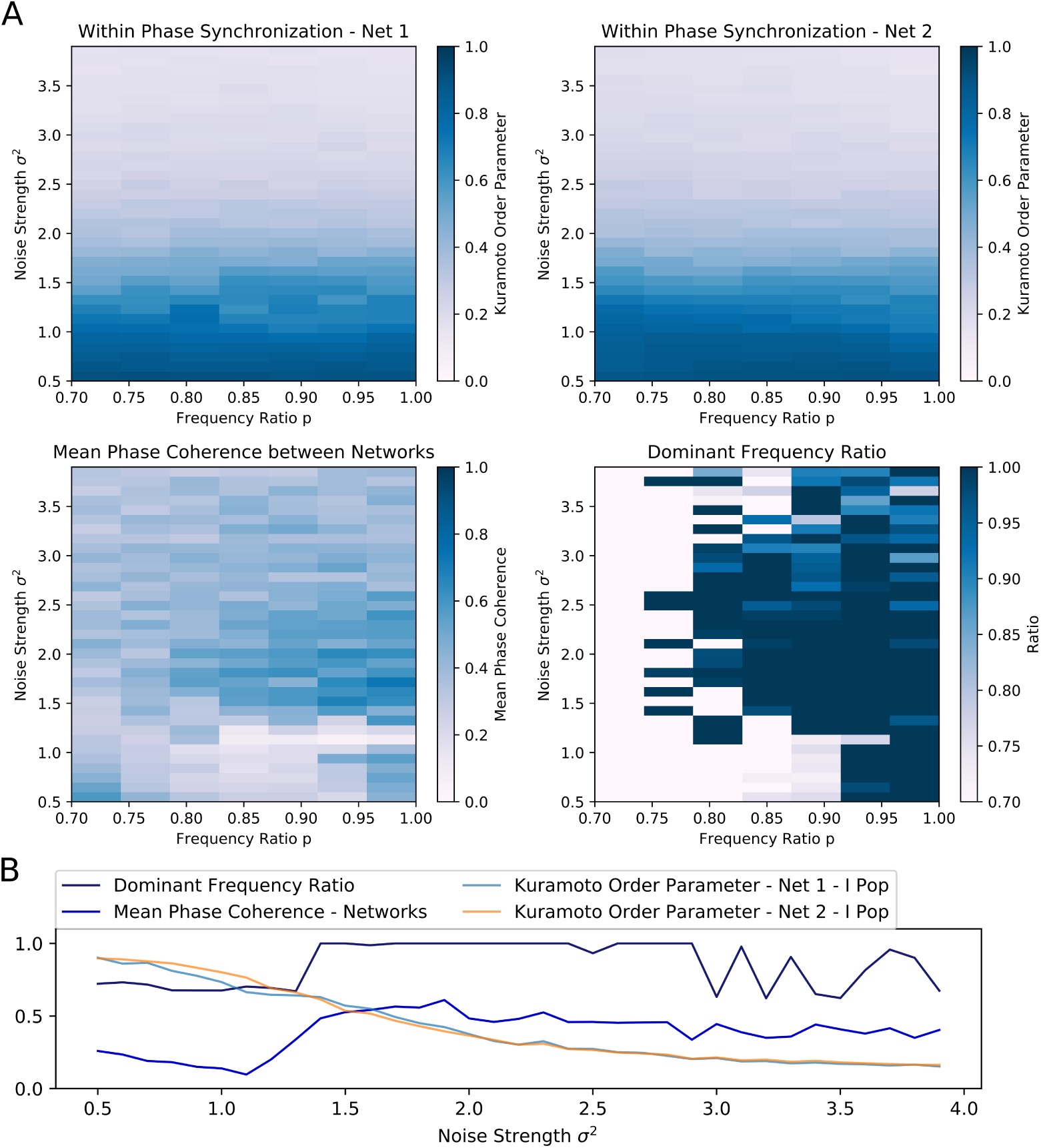
Scenario 1 - Exploration of two interacting all-to-all connected inhibitory networks driven by the ING mechanism. (A) Heat maps visualizing two-dimensional exploration over noise strength *σ*^2^ and noise frequency ratio *p*. The heat maps in the first row encode the phase synchronization within the respective inhibitory network while the bottom row displays phase synchronization and dominant frequency ratio across the two networks. (B) Visualization of a one-dimensional exploration over noise strength *σ*^2^ values from 0.5 to 4.0 in 0.1 steps. The frequency ratio of noise was set to *p* = 0.85.

Furthermore, the results of the one-dimensional exploration (Fig 2B) highlighted the opposing relationship between within and across network synchronization when noise strength *σ*^2^ was increased. In this case, the frequency ratio was initially set to *p* = 0.85.

Again, the dominant frequency ratio and the mean phase coherence between the networks increased while synchronization within the networks decreased. Interestingly, once *σ*^2^ reached 1.4, the network abruptly transitioned to a frequency locked state as the frequency ratio jumped to and stayed at 1.0 while phase locking was increased only slowly with further increase in strength. Although an increase in noise could apparently improve inter-network synchronization, the beneficial effect was limited. In this case, the maximum was reached at *σ*^2^ of 1.9 and a further increase worsened both within and across network synchronization. Additionally, the dominant frequency ratio began to fluctuate at a noise strength of 3.0.

#### Analysis of three distinct states

We then proceeded to compare three parameter configurations, each of which showed a different state in the scenario of two coupled inhibitory networks:

**State 1 Unsynchronized activity across and synchronized activity within networks** - Weak noise and weak inter-network coupling (Fig 3A)

**Fig 3.**
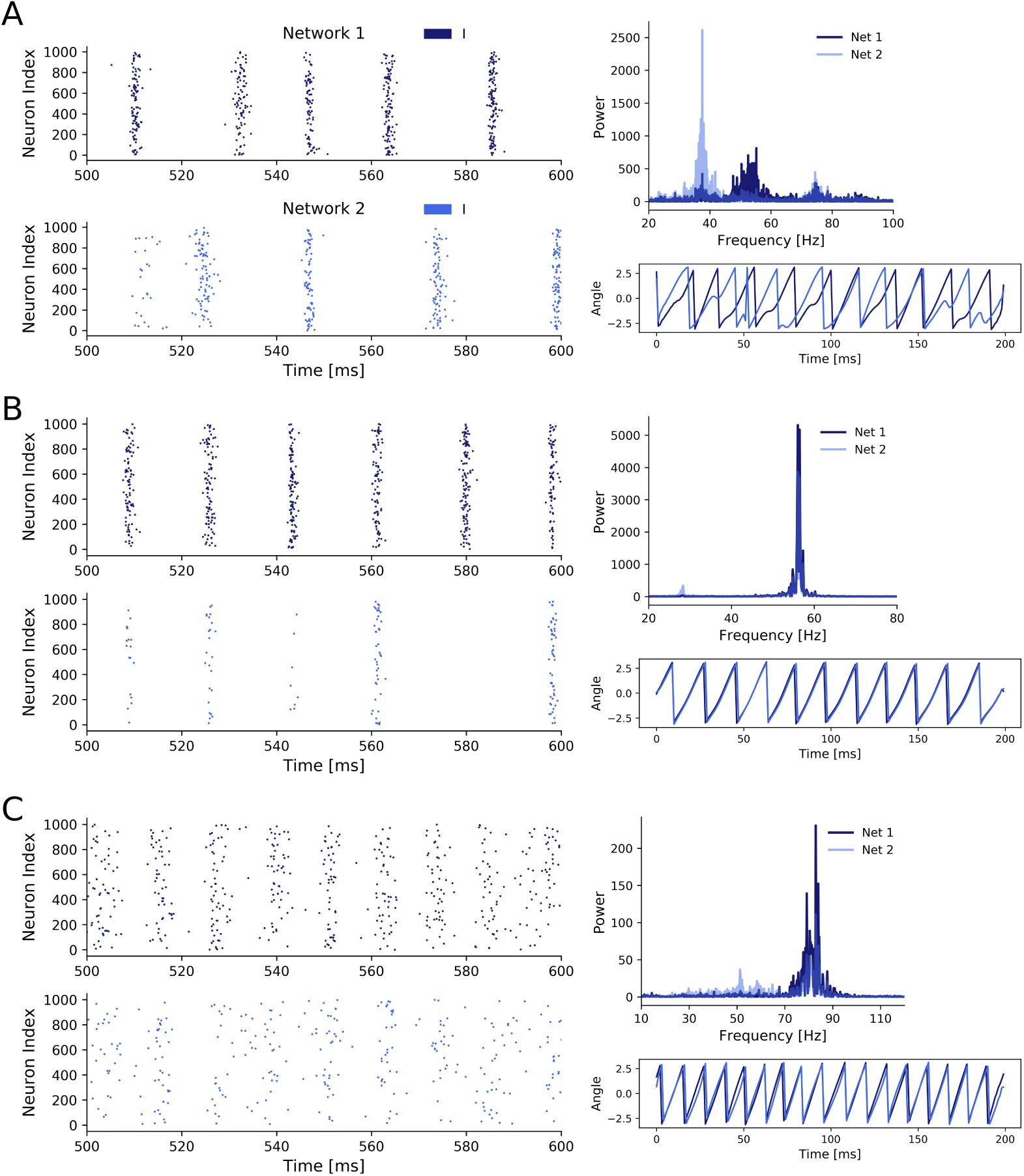
Three states of scenario 1. (A) Two coupled ING networks subject to weak independent noise and low inter-network coupling. With *p* = 0.85, *σ*^2^ = 0.5, *J*_*ppii*_ = 0.15. (B) ING networks subject to weak independent noise and strong inter-network coupling. With *p* = 0.85, *σ*^2^ = 0.5, *J*_*ppii*_ = 0.3. (C) ING networks subject to strong independent noise and weak inter-network coupling. With *p* = 0.85, *σ*^2^ = 1.5, *J*_*ppii*_ = 0.15.

**State 2 Synchronized activity across and within networks** - Weak noise and strong inter-network coupling (Fig 3B)

**State 3 Synchronized activity across and unsynchronized activity within networks** - Strong noise and weak inter-network coupling (Fig 3C). In contrast to the second state, synchronization within the networks was impaired, pointing towards a different synchronization mechanism.

Additionally to the case of weak inter-network coupling with weak and strong noise (state 1 and 3, respectively), which was already explored in Meng and Riecke [32], we here also included the case of strong inter-network coupling with weak noise. As we will demonstrate later, state 2 and 3 both displayed synchronization across networks, however the underlying synchronization mechanism was fundamentally different. Thus, we here extend the findings of Meng and Riecke [32] for ING networks.

State 1 (Fig 3A) was characterized by a high within-network synchronization and no synchronization across networks. This was not surprising, since the coupling strength between networks and the noise strength were both comparatively weak so that the behaviour of one network was not considerably influenced by activity from the respective other network. Specifically, there was a mismatch between the dominant frequencies of the two networks (55 Hz for network 1 and 39 Hz for network 2), because network 2 received noisy spike input with a lower rate determined by the noise frequency ratio *p*, in this case *p* = 0.85. Unsurprisingly, the temporal evolution of the networks’ phases also did not display any coherence between the two networks. Further, the coherent and rhythmic firing of the inhibitory neurons confirmed the high phase synchronization within each network.

For state 2 (Fig 3B), coupling strength between the networks was increased until the two networks synchronized. This state was characterized by a homogeneous behaviour within networks *and* across networks, demonstrated by the overlapping power spectra and matching dominant frequencies, confirming a 1:1 frequency locking. This was further confirmed by the evolution of the phases of the networks. Interestingly, while both networks again displayed rhythmic behavior, the participation of neurons in network 2 was markedly reduced compared to network 1 and only a small fraction of the neurons participated in each cycle. The networks synchronized through a winner-takes-all effect in favour of the faster network 1. By activating inhibitory neurons in network 2, network 1 suppressed all neurons in network 2 that did not spike precisely in cycles of the inhibitory rhythm of network 1. Over time, the population activity of network 2 synchronized with the population activity of network 1. Again, the total power in network 2 was lower compared to network 1, since network 2 received a lower proportion of the noise input and was subject to strong inhibition from the faster network 1.

In state 3 (Fig 3C), when we increased noise strength while keeping inter-network coupling weak, the networks also transitioned to a 1:1 phase and frequency locked state. But contrary to state 2, this was achieved through the increased variability in the membrane potentials, which weakened the synchronization within a network, while allowing a variable fraction of neurons in the slower network 2 to participate in cycles of network 1. The increase in noise strength increased spike time variability inside a network, thereby weakening the gating effect of inhibition, which led to an acceleration of population rhythms and explained the higher dominant frequencies in the power spectra of both networks compared to state 1 and 2. Similar to state 2, the dominant frequencies matched, but the noise led to a spread in the power of both network signals, especially in network 2. Furthermore, the networks were in general 1:1 phase locked despite some occasional irregularities caused by the higher variability of spiking in both networks due to the strong noise. In state 2, strong inhibition from network 1 led to suppression of spikes in network 2 if they did not fall into cycles of network 1, explaining their synchronization over time. However, in state 3 synchronization was induced by a different mechanism. Specifically, the strong noise weakened the within-network synchronization (especially in the inhibitory population) as inhibitory feedback was not able to completely cancel out the noise-induced randomness contrary to the weak noise in state 2. This reduced inhibition inside a network enabled spikes of neurons in network 1 to have a bigger influence on the neurons in network 2, explaining the sparse spike participation of network 2 in cycles of network 1.

### Uncorrelated noise can synchronize networks of PING populations

Next, we explored the effects of uncorrelated noise in interconnected networks of excitatory and inhibitory populations. We investigated two types of EI networks, all-to-all coupled (scenario 2) and random, sparsely coupled networks (scenario 3). However, since the findings were independent of the particular coupling type we only present the results for the biologically more plausible scenario of sparse and random connectivity between neurons in each population. The results for the all-to-all coupled networks can be found in the Supplementary Material.

#### Exploration

In the case of two random sparsely connected excitatory-inhibitory networks, increased noise strength could indeed synchronize the two networks with respect to phase and frequency locking (Fig 4A). Similar to scenario 1, for a fixed frequency ratio *p*, increased noise strength *σ*^2^ eventually pushed the unsynchronized networks to a synchronized state. Furthermore, there was again an inverse relationship between the frequency ratio *p* between the networks and the noise strength required for the networks to synchronize their rhythms.

**Fig 4.**
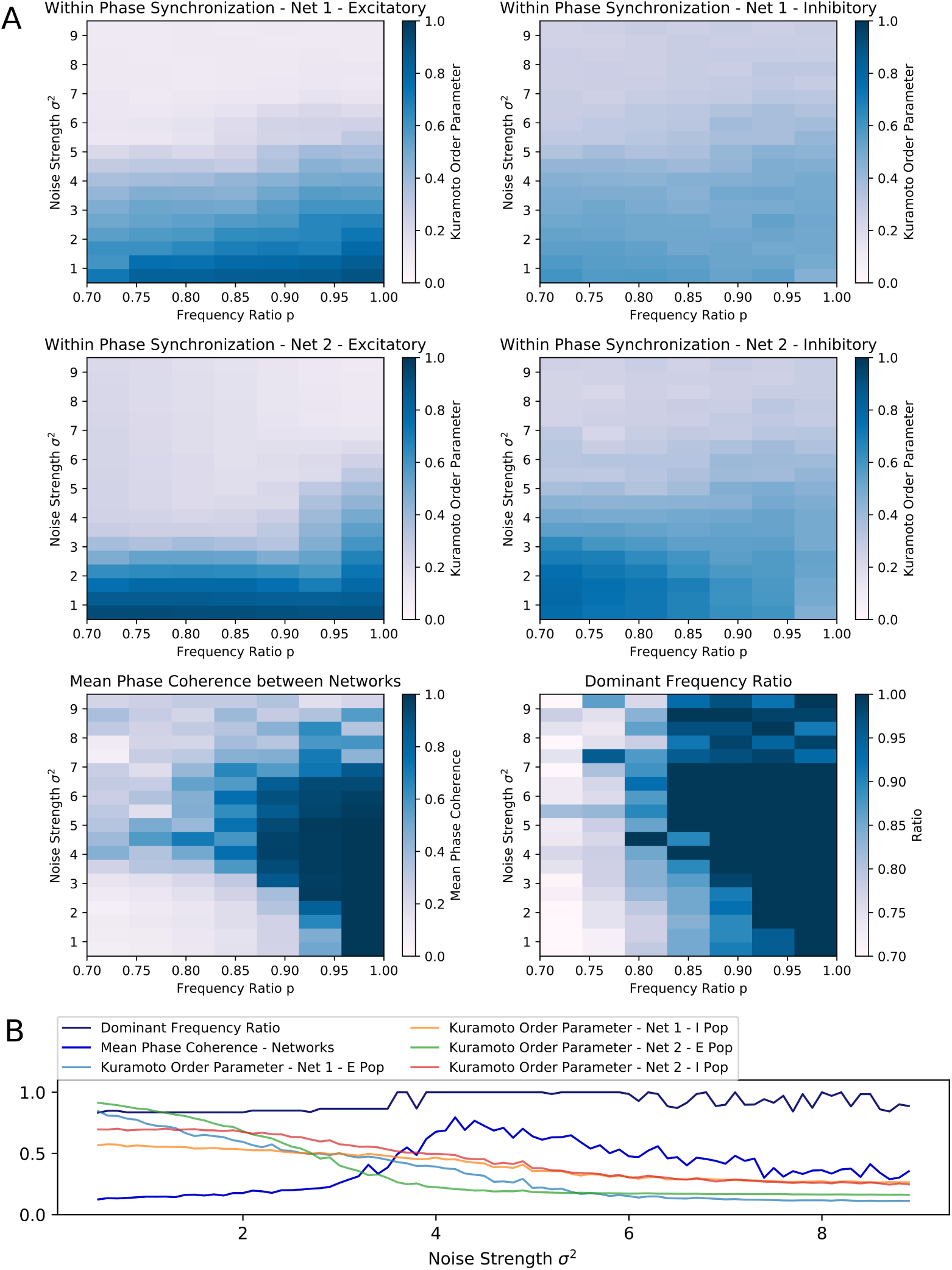
Scenario 3: Exploration of two random sparsely connected excitatory-inhibitory networks driven by the PING mechanism. (A) Exploring the within and across network synchronization behavior over different noise strengths *σ*^2^ and noise frequency ratio *p* values. (B) One-dimensional explorations over noise strength *σ*^2^. Noise frequency ratio stayed constant with *p* = 0.85. Range of 0.5 to 9.0 in 0.1 steps with runtime of 3s for each trial.

Interestingly, the PING rhythms seemed to be more robust against noise than the ING rhythms in scenario 1. It required comparatively higher noise strengths to produce any noticeable changes in the behaviour of each network. However, as soon as the noise strength was increased sufficiently, high values of the mean phase coherence and the dominant frequency ratio between the two networks could be observed. Again, this beneficial effect was bounded from above and further increases in noise strengths led to a deterioration of both within *and* across network synchronization, suggesting that network behavior was mainly determined by the external noise input in this parameter regime.

The parameter space in which frequency and phase locking could be observed was reduced and limited to input frequency ratios above 0.80 (Fig 4A). The sensitivity to noise was also visible based on the fast decline of within phase synchronization in both networks when noise strength was increased independently of the frequency ratio p. Again, the beneficial effect of noise was bounded and noise strengths above 7 led to low synchrony within *and* across networks (Fig 4A).

Furthermore, the results of the one-dimensional exploration of scenario 3 confirmed an inverse relationship between inter-network synchronization measures and within-network synchronization measures over a wide range of noise strengths (Fig 4B), matching the results from scenario 1. Together with the mean phase coherence the dominant frequency ratio reached high values at a certain strength threshold (at approx. *σ*^2^ = 3.7), while the Kuramoto order parameter of each population decreased nearly monotonically with increasing noise strength.

Again, depending on the noise frequency ratio, increasing the noise beyond a certain threshold had a detrimental effect on all synchronization measures and eventually led to fluctuating dominant frequency ratios (Fig 4B).

#### Analysis of three distinct states

We again investigated the same three different states as for the ING networks (see previous section):

**State 1 Unsynchronized activity across and synchronized activity within networks** - Weak noise and weak inter-network coupling (Fig 5A)

**Fig 5.**
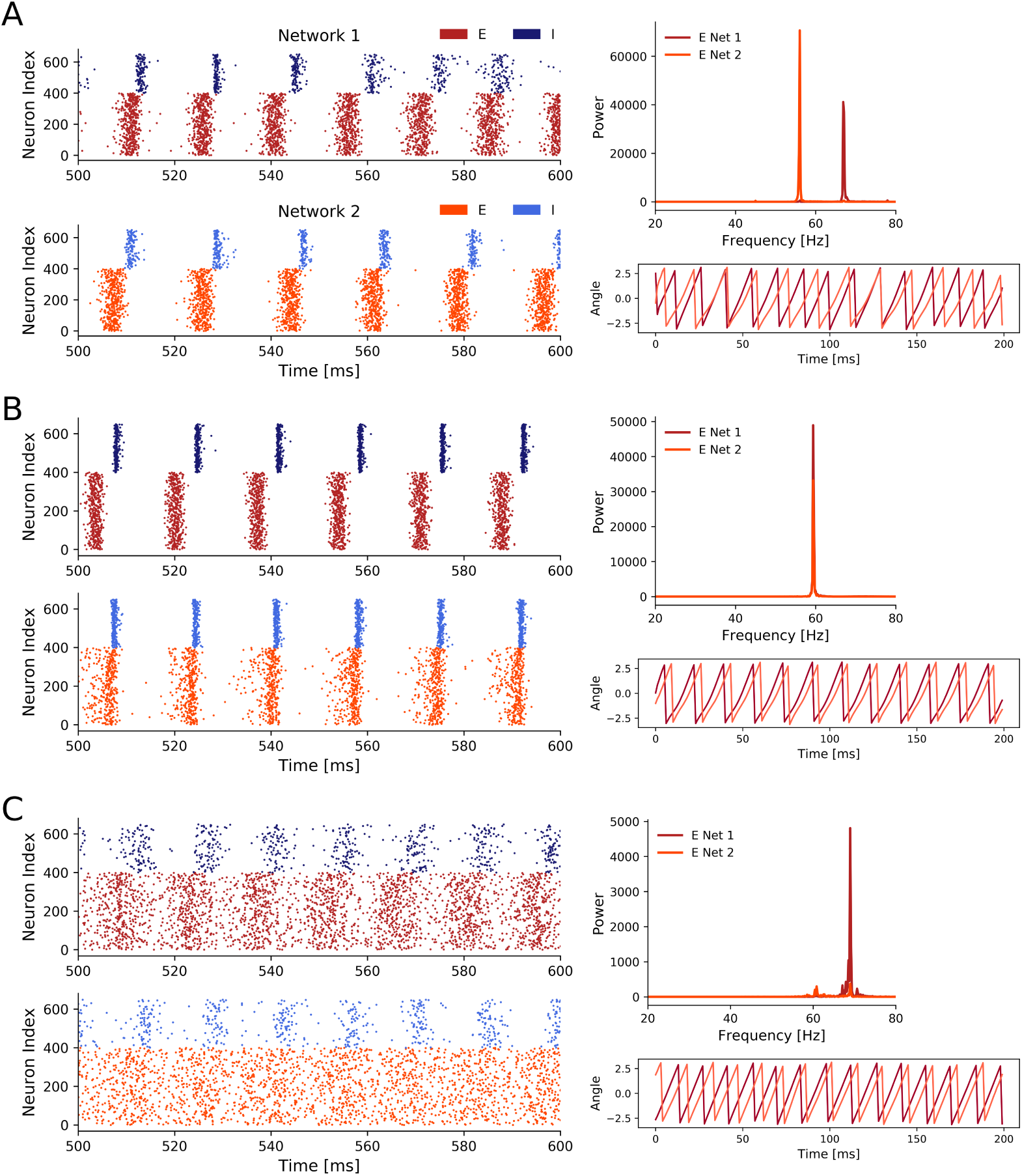
Three representative states of scenario 3. (A) Weak noise and weak inter-network coupling. With *p* = 0.85, *σ*^2^ = 0.7, *J*_*ppei*_ = 0.03. (B) Coupling strength was increased until we observed 1:1 frequency entrainment. With *p* = 0.85, *σ*^2^ = 0.7, *J*_*ppei*_ = 0.07. (C) Strong noise and weak inter-network coupling. With *p* = 0.85, *σ*^2^ = 4.5, *J*_*ppei*_ = 0.03. Only 400 out of 1000 excitatory (red) neurons are displayed in the spike raster plots to reduce plot size.

**State 2 Synchronized activity across and within networks** - Weak noise and strong inter-network coupling (Fig 5B)

**State 3 Synchronized activity across and unsynchronized activity within networks** - Strong noise and weak inter-network coupling (Fig 5C).

Overall, for all three states we saw clear PING rhythms in both EI networks, where excitatory neurons fired first driving the inhibitory population which subsequently silenced the excitatory activity. After the decay of the inhibition the excitatory neurons fired again and the gamma cycle started over. However, the participation of neurons in each gamma cycle varied for the three states. While it was high for states 1 and 2, it was lower for the third state with strong noise and weak inter-network coupling.

In state 1 (Fig 5A), the low inter-network coupling combined with the weak noise input was not sufficient to synchronize the two networks. Therefore, the dominant frequencies of both networks were solely determined by the frequency ratio *p* = 0.85, resulting in a faster rhythm at 68 Hz for network 1 (which received stronger noise) and a slower rhythm at 58 Hz for network 2. As the rhythms between the two networks were not synchronized, the phases of the networks showed, unsurprisingly, no coherence.

Once we increased the inter-network coupling strength sufficiently in state 2, the networks synchronized their rhythms (Fig 5B). This was expressed by a match of the dominant frequencies at 60 Hz. Noteworthy, the increased coupling slowed down the rhythm of network 1. The phase plot showed a phase locked state where the phases display a constant difference with each other. Both networks displayed a strong PING rhythm and the increased coupling between the networks, compared to state 1, reduced the variability in the spike times of the I populations.

In state 3 (Fig 5C), while an increase in the noise strength *σ*^2^ transitioned the networks to a state in which their oscillations were frequency and phase locked, similar to an increase in the inter-network coupling strengths, the synchronization within these networks was considerably decreased which was shown by the high spike variability in the spike raster plots. Interestingly, in contrast to state 2 where the rhythm of network 1 slowed down to the pace of network 2, we observed the opposite behavior in state 3. The slower network 2 sped up and both networks shared a peak activity at 68-70 Hz. Compared to state 1, in state 3 phase synchronization was increased, although it did not reach the nearly perfect synchronization achieved in state 2. Although the PING rhythm was still present in both networks, variability in the firing of inhibitory as well as excitatory cells was markedly increased by the strong uncorrelated noise. Especially the firing of the E cells was spread widely across the time interval between two consecutive inhibitory firing cycles. This explained the distinct decrease of the power in the frequency spectra compared to state 1 and 2.

### Details of the synchronization mechanism

After demonstrating that strong, uncorrelated was also able to synchronize interconnected EI networks, we looked at the synchronization mechanism in more detail. Again, the mechanism is general to the all-to-all (scenario 2) and sparse random connectivity (scenario 3), so we limited our analysis to the latter to minimize redundancy. Based on Meng and Riecke [32] and our previous analysis of two interacting inhibitory networks (scenario 1), we expected that the increased heterogeneity in the inhibitory membrane potentials caused by sufficiently strong uncorrelated noise would also be a key factor in the noise-induced synchronization between EI networks.

#### Heterogeneity in inhibitory membrane potentials

As is evident from the previous section, the synchronization induced by strong inter-network coupling in state 2 was fundamentally different from synchronization induced by an increase in the noise strengths present in state 3. Specifically, in state 3 the inhibitory cells of the second network can be divided into two groups based on an arbitrary gamma cycle *c*_*i*_ of the inhibitory population in the first network (Fig 6A). The first group fired in cycle *c*_*i*_ in response to the firing of the excitatory population in network 1. However, the neurons in the second group stayed suppressed and skipped this cycle. However, they were more likely to fire in the next cycle *c*_*i*+1_. Thus, there was always a fraction of neurons that was likely to fire at a cycle *c*_*i*_ in response to excitation from network 1 and a remaining fraction that already fired in one of the previous cycles *c*_*i−*1_, *c*_*i−*2_, … and therefore skipped cycle *c*_*i*_, in order to fire with increased probability in one of the upcoming cycles *c*_*i*+1_, *c*_*i*+2_, ….

**Fig 6.**
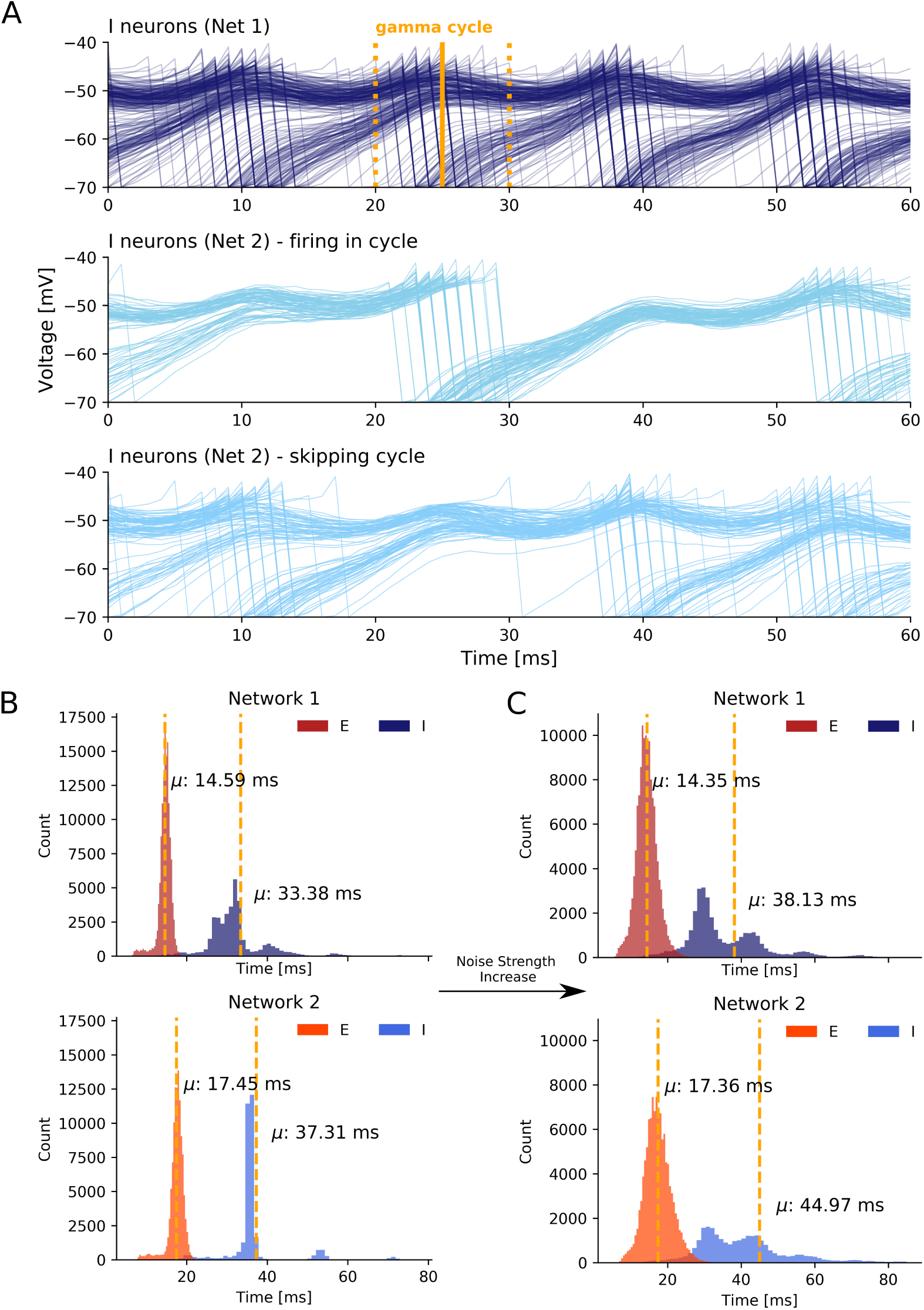
Synchronization mechanism based on uncorrelated noise. (A) The first plot displays voltage traces of I neurons in network 1 while the second and third plot display grouped voltage traces of I neurons in network 2. We selected an arbitrary I cycle *c*_*i*_ of network 1 in state 3 marked by the time window [*t*_*start*_, *t*_*end*_] (yellow lines). A fraction of neurons participated in the selected cycle (second plot) while the remaining neurons skipped the cycle and sparsely participated in the previous and next cycle (third plot). The values on the x axis are relative. (B) ISI histogram of state 1 with weak coupling and weak noise. (C) ISI histogram of state 3 with weak coupling and strong noise.

Overall, the noise-induced variability of the membrane potentials weakened the PING rhythm within the networks and thereby enhanced the responsiveness of neurons to excitation from the respective other network outside of the temporal window of the neurons’ network rhythm. This caused the slower network 2 to speed up to the pace of the faster network 1.

#### Interspike interval histograms

In order to quantify the observed spike variability, we next took a look the interspike interval (ISI) histograms for each population. To get insight into how stronger noise transitions the weak synchrony of state 1 into the high synchrony state 3, we examined the ISI histograms of the I and E cells of both networks (Fig 6B and C). For state 1, we observed relatively low variance in all four groups. Importantly, with strongly increased noise the model transitioned from state 1 to state 3 and the variance of the ISIs of both the E and, even more pronounced, the I populations was noticeably increased in state 3 (Fig 6C). Besides an increase in the variance, the mean ISIs *μ* increased in both I populations as well. As the dominant frequency in network 1 did not change from state 1 to state 3 and as the dominant frequency in network 2 even increased in state 3, the higher mean ISI implied a decrease in the participation of I cells in their respective population rhythm. Interestingly, the ISIs of E cells did not change as much as those of the I cells over the transition from state 1 to state 3, although the E populations received a higher amount of noise as input (we increased *σ*^2^ from 0.7 to 4.5). Conclusively, the increase in the ISI variability and mean of the I populations confirmed the current notion that the noise-induced heterogeneity of the I cells is a key factor in enhancing synchronization between EI networks.

## Discussion

We explored the role of uncorrelated noise in the synchronization of interacting gamma rhythms. We confirmed prior results from Meng and Riecke [32] on ING rhythms and extended their findings to gamma oscillations produced by the PING mechanism. To this end, we modelled two interconnected excitatory-inhibitory networks in various network settings and analyzed how synchronization within and across the networks changed depending on the noise strength.

### Uncorrelated Noise facilitates Inter-Network Synchronization in PING Networks

We found an optimal range of noise strengths that enhanced across network synchronization. However, the beneficial effect of noise was limited and we detected an upper and lower bound in all three scenarios. Noise strengths above the upper bound eventually broke the network rhythms as spiking of neurons was mainly determined by the uncorrelated input, indicated by a worsening of within *and* across network phase synchronization. On the other hand, noise strengths below the lower bound were not able to dampen the within network coupling sufficiently to enhance the responsiveness of neurons to input from the respective other network.

### Noise-induced Synchronization Mechanism

Further, we confirmed that the mechanism described by Meng and Riecke [32] that underlies the noise-induced synchronization across networks in inhibitory networks was also present in the PING networks considered here. Although the two investigated variants of two interacting EI networks differed in several points from inhibitory networks, the same fundamental mechanism could be observed. Noise caused considerable voltage fluctuations in all neurons, thereby weakening the PING rhythm within a network. This led to an increased responsiveness of neurons in one network to excitation from the respective other network as a fraction of neurons was likely to be close to its membrane threshold. This enabled a variable fraction of neurons to spike in response to excitation from the other network, thereby promoting the network rhythms to synchronize over time with the activity of the faster network. Noteworthy, in the case of inhibitory networks, it was not *excitation* of the faster network that sped up the rhythm of the slower network. Instead, *inhibition* of the faster network gated the slower network similar to the ING mechanism itself [32].

Importantly, the identified synchronization mechanism is fundamentally different from *stochastic synchronization* promoted by correlations in the external noise input (as already discussed in [32]). Further, it also differs from *stochastic resonance* [39, 40] *or the related phenomenon of enhanced responsiveness* [41] where noise amplifies a weak input stimulus so that it can be detected by the neuron. While the identified mechanism enhanced the responsiveness of neurons to excitation from the other network, it did not amplify a weak signal but instead weakened the rhythm within a network. This rendered neurons susceptible to spikes in response to excitation or inhibition outside of the temporal window defined by the networks own rhythm. We also showed that the noise-induced synchronization is different from synchronization introduced through strong coupling between the networks and that they show different signatures. While coupling-induced synchronization leads to a high within- and between-network synchronization, noise-induced synchronization shows weaker within-network synchronization together with high between-network synchronization. Furthermore, noise-induced synchronization led to a speed up of the slower network to match the oscillation frequency of the faster network, whereas synchronization through strong coupling resulted in a slowing down of the fast network. We propose the following view: the fundamental aspect of the noise-induced synchronization mechanism is the desynchronization within a network that enhances the responsiveness of neurons to *any external stimulation outside of the short temporal window* defined by the ING or PING mechanism, in this case a second network with a faster rhythm.

As the spiking behavior of the E population did not considerably vary across scenarios, we hypothesize that noise-induced variability in the inhibitory population is central to enhancing synchronization between networks and that the desynchronization of the E population has a supporting role by facilitating the heterogeneity of the inhibitory population.

### Role of Uncorrelated Noise in Aberrant Neuronal Communication

Importantly, the communication through coherence (CTC) hypothesis suggests that neuronal communication among neuronal groups is mediated by phase synchronization [17, 19]. Conclusively, our findings suggest that uncorrelated noise, mimicking the strong synaptic noise observed in the cerebral cortex [42], can have a supporting role in facilitating neuronal communication among neuronal networks displaying rhythmic gamma band activity. In specific, there exists an optimal level of noise that allows the transition of networks from a desynchronized state to a synchronized state by enhancing the responsiveness of neurons inside a network to inhibitory and excitatory input originating from another network. This suggests that deficits in sensory or cognitive abilities as seen in several neurological or psychiatric disorders which have been hypothesized to be related to aberrant synchronization in the gamma band, might be due to an increase in the signal-to-noise ratio that is often seen in these disorders. For example, patients with schizophrenia show deficits in visual Gestalt perception [43] and working memory performance [44], two cognitive processes that have been linked to gamma oscillations. Furthermore, it has been shown that the signal-to-noise ratio in neural activity of patients with schizophrenia is significantly increased [45–48]. Therefore, these deficits might be attributable to a decreased ability of inter-network synchronizability in the gamma band due to the increased noise levels.

### Limitations

The model we employed in our explorations is of course a highly simplified representation of neural populations *in vivo*. For instance, our model currently represents a standard EI model of one excitatory and one inhibitory population and ignores the existence of multiple inhibitory populations of different interneuron types. We only modelled fast-spiking parvalbumin-expressing (PV^+^) interneurons since their somatic inhibition of regulates the timing of action potentials and imposes brief time windows in which PCs can spike, therefore, promoting synchronization [49]. Furthermore, PV^+^ interneurons seem to be responsible for the generation of gamma oscillations [50–52]. However, somatostatin-expressing (SST^+^) and vasoactive peptide-expressing (VIP^+^) interneurons contribute substantially to regulating GABAergic inhibition in the cortex [53–55] and dysfunction of these interneurons are associated with psychiatric disorders [55, 56]. Additionally, SST^+^ interneurons are presumed to play an important role in synchronization of visually induced, context-dependent gamma rhythms in visual cortex [54].

Additionally, our model does not represent the layered structure of cortex and therefore cannot capture the intricate differences in information processing between different layers of cortical regions and their oscillatory signatures.

Furthermore, in our current model we simply feed external, uncorrelated noise into the two networks and do not model the origin of this noise. Further studies are warranted to elucidate the effect of different sources of uncorrelated noise in cortical networks, such as stochastic synaptic transmission and ion-channel noise [57, 58], on the synchronization mechanism described here. We also did not study the interaction this noise-induced synchronization mechanism with other synchronization mechanisms due to correlated noise, such as stochastic synchronization [59–62], that have been described before.

## Conclusion

We replicated the findings of Meng and Riecke [32] in two coupled inhibitory networks of AdEx neurons that display gamma band activity produced by the ING mechanism. Next, we extended their work to excitatory-inhibitory coupled networks displaying gamma rhythms produced by the PING mechanism. In specific, our results show that certain levels of uncorrelated noise can enhance the synchronization *across* interacting EI networks although worsening the synchronization *within* these networks.

Further, we confirmed the underlying mechanism described by Meng and Riecke [32]. It is the noise-induced variability that weakens the rhythm within a network and thereby enhances the responsiveness of neurons to input from respective other networks, eventually resulting in the synchronization of the network rhythms.

In light of the CTC hypothesis [17, 19], our work suggests that the strong synaptic noise observed in the cerebral cortex can have a supporting role in facilitating neuronal communication among neuronal networks displaying gamma band activity. We hypothesize that coherence of gamma band activity between interacting neuronal groups could be enhanced by raising or lowering synaptic noise in at least one of the groups.

## Supplementary Material

**Fig 7.**
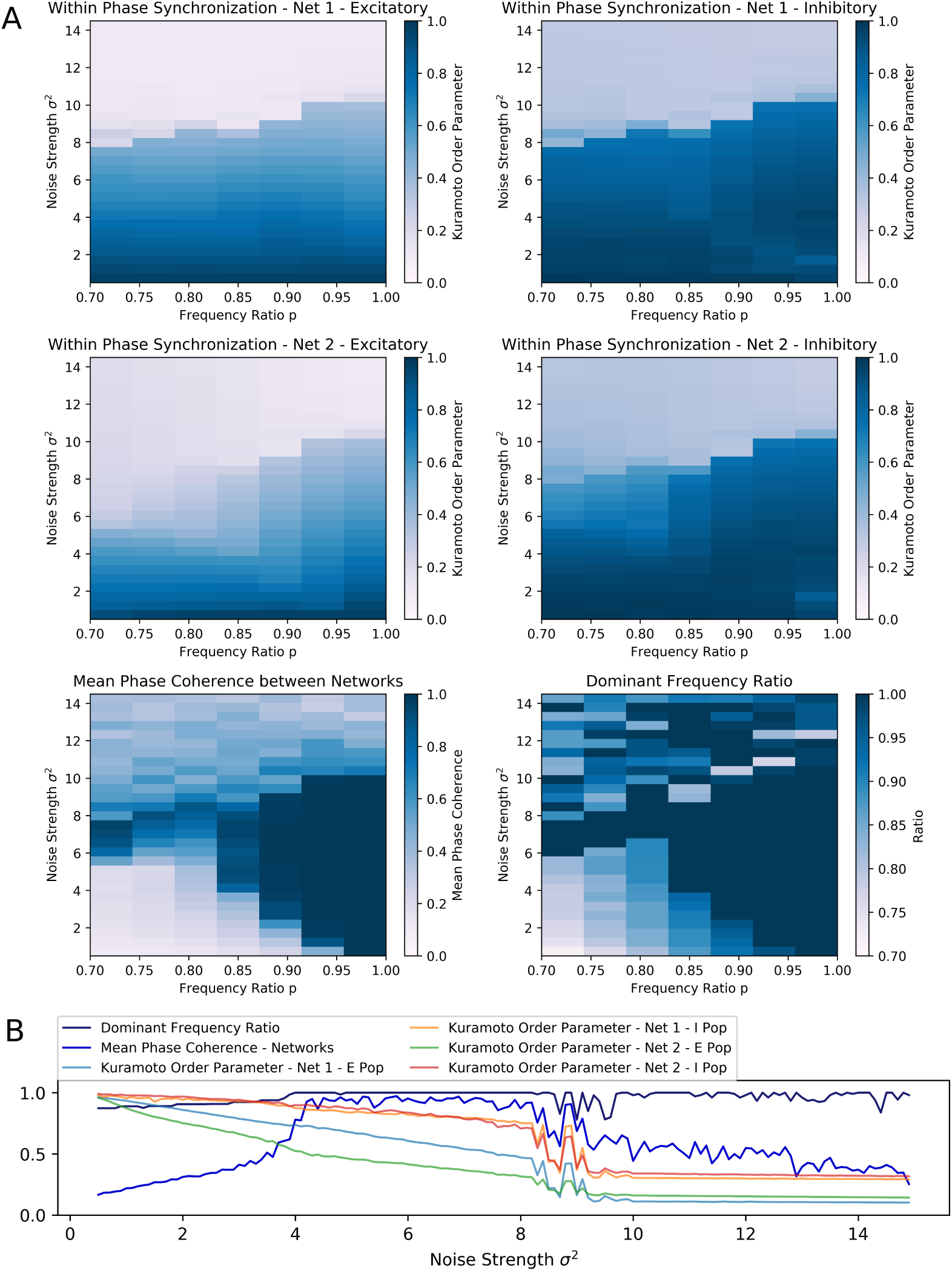
Scenario 2: Exploration of two interacting all-to-all connected excitatory-inhibitory networks driven by the PING mechanism. (A) Exploring the within and across network synchronization behavior over different noise strengths *σ*^2^ and noise frequency ratio *p* values. (B) One-dimensional explorations over noise strength *σ*^2^. Noise frequency ratio stayed constant with *p* = 0.85. Range of 0.5 to 15.0 in 0.1 steps with runtime of 3s for each trial.

## Footnotes

1 https://www.scipy.org/

2 https://matplotlib.org/

3 https://github.com/ChristophMetzner/Synchronization-by-Uncorrelated-Noise

4 https://pypi.org/project/matplotlib/

5 https://www.scipy.org/

6 https://brian2.readthedocs.io/en/stable/user/numerical_integration.html

